# N-cadherin mechanosensing in ovarian follicles controls oocyte maturation and ovulation

**DOI:** 10.1101/2023.10.06.561232

**Authors:** Alaknanda Emery, Orest W. Blaschuk, Thao D. Dinh, Tim McPhee, Rouven Becker, Andrew D. Abell, Krzysztof M. Mrozik, Andrew C.W. Zannettino, Rebecca L Robker, Darryl L Russell

## Abstract

The cell adhesion molecule N-cadherin (CDH2) is a membrane component of adherens junctions which regulates tissue morphogenesis and architecture. In the follicles of mammalian ovaries, N-cadherin adherens junctions are present between granulosa cells, cumulus cells and at the interface of cumulus cell transzonal projections and the oocyte. We demonstrate a mechanosensory role of N-cadherin integrating tissue structure and hormonal regulation of follicular morphogenic events including expansion of the cumulus oocyte complex (COC) matrix, oocyte maturation and ovulation. Two small molecule N-cadherin antagonists inhibited COC maturation *in vitro*. Transcriptome profiling revealed that targets of β-catenin and YAP1 pathways were dysregulated by N-cadherin antagonists. *In vivo*, N-cadherin antagonist significantly reduced ovulation in mice compared to controls (11 vs 26 oocytes/ovary; p=5.8×10^-6^). Ovarian follicles exhibited structural dysgenesis with granulosa and cumulus cell layers becoming disorganised and the connection between cumulus cells and the oocyte disrupted and the transcriptome again indicated altered mechanical sensing causing dysregulation of the Hippo/YAP and β-catenin pathways and ECM reorganization. Granulosa specific N-cadherin depletion in Cdh2^Fl/FL^;Amhr2-Cre also showed significantly altered mechanosensitive gene expression and reduced ovulation. Our findings demonstrate a critical role for N-cadherin in ovarian follicular development and ovulation, and the potential to inhibit ovulation through targeting this signalling mechanism.

## INTRODUCTION

Classical (type I) cadherins are calcium dependent cell adhesion molecules that form intercellular adherens junctions thereby regulating cell-cell recognition and the assembly of tissues. These cadherins control a myriad of biological processes including cell migration, proliferation, death and adhesion [1, 2]. They consists of extracellular, transmembrane and intracellular domains. The intracellular domain interacts with a complex composed of β-catenin, α-catenin, p120 and α-actinin that links the Type I cadherins to the actin cytoskeleton [3, 4]. The formation of adherens junctions requires interaction of cadherins within the same membrane to form lateral or *cis*-dimers. These *cis*-dimers then promote intercellular adhesive dimerization or *trans*-dimers of cadherins on adjacent cells [5] to form adherens junctions. Intracellular signalling pathways are activated by mechanical force generated through cell-cell and cell-matrix interactions. Two types of cell surface receptors, cadherins and integrins directly mediate cell adhesion and activate intracellular pathways such as β-catenin [6]. Cross-talk between the classical cadherin, N-cadherin (CDH2) and the Hippo signalling pathway is another mechanosensitive signalling axis that regulates cell proliferation and determines organ size [7]. The Hippo-YAP1 signalling is regulated by numerous upstream signals including organ size, extracellular matrix (ECM) stiffness and cell polarity [8].

Ovarian follicles are complex multilayered structures required for the development and maturation of oocytes. Follicles also serve as an endocrine organ coordinating the female reproductive cycle, augmenting oocyte maturation and ovulation (the release of the oocyte) at the appropriate time for fertilization. In growing follicles, the cumulus cells surrounding the oocyte provide essential metabolites and maintain oocyte meiotic arrest through transfer of cAMP and cGMP from cumulus cells to oocyte. This communication requires thin cytoplasmic projections known as transzonal projections (TZPs) that extend from the cumulus cells, through the thick ECM of the zona pellucida to contact the oocyte plasma membrane [9]. Gap junctions at the cumulus-oocyte cellular contacts transport small molecules such as glucose metabolites, ATP and cAMP between the cells [10]. Adherens junctions have been proposed to stabilise these cumulus cell-oocyte contacts as a necessary precursor to gap junction formation [11], and the contacts are selectively uncoupled just prior to ovulation [12].

Contacts between cumulus and granulosa cell layers are critical for the progressive development of ovarian follicles and maturation of oocytes [9, 13, 14]. N-cadherin is present on the intercellular junctions between granulosa cells [15] and β-catenin has been observed on granulosa and cumulus cell membranes, while another Type I cadherin, E-cadherin (CDH1) is expressed on oocyte membranes [16, 17]. β-catenin is also a transcription factor which plays an important role during follicle development and granulosa cell specification in the ovary [18]. Constitutive activation of β-catenin in Sertoli cells of the mouse testis drives their trans-differentiation toward ovarian granulosa cell lineage [19], while in ovaries constitutive β-catenin induces elevated expression of the estrogen synthesising gene *Cyp19a1* [20]. YAP1 also plays a critical role in mediating regulation of follicle development through mechanical cues [21]. In granulosa cells YAP1 is required for proliferation and promotes differentiation [22]. However, the respective roles of N-and E-cadherin regulation of β-catenin and YAP1 pathways during ovarian folliculogenesis or oocyte maturation and ovulation has not been investigated.

Ovulation is initiated by a surge of luteinizing hormone (LH) from the pituitary, inducing dramatic remodelling of the ovarian follicle ECM, and differentiation of cells. The intercellular junctions between cumulus cells and the oocyte are disrupted as the COC undergoes dynamic expansion of a unique viscoelastic ECM through gene induction in granulosa and cumulus cells [23, 24]. This COC matrix expansion is essential for ovulation and fertilization-potential of oocytes [23, 25]. The expanded COC acquires an adherent and migratory cell phenotype and can be shown to robustly adhere to ECM components such as fibronectin and collagens that are abundant in the follicle wall [26].

The localisation of N-cadherin on granulosa cells was recently shown to be involved in primordial follicle formation (reviewed in [27]; [28]), and N-cadherin antagonists have also been shown to cause apoptosis of cultured granulosa cells [29]. We now report previously undiscovered roles for N-cadherin in granulosa cell responses to hormones, as well as COC expansion, oocyte meiotic maturation, and ovulation. Two recently developed, small molecule N-cadherin antagonists [4, 30] were shown to disrupt the interaction between TZPs and oocytes, and to dysregulate β-catenin and YAP1 signalling pathways in mouse COCs, blocking oocyte meiotic maturation. Furthermore, treatment of mice *in vivo* with an N-cadherin antagonist blocked ovulation with pervasive effects on β-catenin and YAP1 pathways leading to altered response to ovulation stimulus. Our findings demonstrate a role for N-cadherin mechanosensing to maintain ovarian tissue architecture, and hence the hormone responsiveness essential for oocyte maturation and ovulation.

## RESULTS

### N-cadherin antagonists block acquired periovulatory COC adhesion

Through screening a small molecule library, we identified two compounds that block the adhesion of periovulatory COCs to fibronectin using the xCELLigence^TM^ system. Two potent COC adhesion blocking compounds were; (S)-1-(3,4-Dichlorophenoxy)-3-(4-((S)-2-hydroxy-3-(4-methoxyphenoxy) propylamino) piperidin-1-yl)propan-2-ol (CRS-066) [30] and 5[3,4 dichlorobenzyl)sulfanyl]4H 1,2,4 triazol 3 amine (LCRF-0006) [4], both of which are small molecule N-cadherin antagonists. Dose-dependent inhibition of COC adhesion to fibronectin, compared to vehicle controls, was confirmed at doses above 1μM for CRS-066 (IC50=0.4μM) and above 36μM for LCRF-0006 (IC50=38 μM) (Fig. 1a). Analogues of both compounds were synthesised with substitutions of two key active chloride side chains on the aromatic ring structure, which abolished the COC adhesion blocking activity at the same concentrations (Fig. 1a). We confirmed the N-cadherin blocking action of the active compounds by showing they reduced N-cadherin abundance at intercellular contacts in N-cadherin expressing SKOV-3 ovarian cancer cell line [31]. Again, significant effects of CRS-066 and LCRF-0006 were observed with 1 μM and 36 μM respectively (Fig. 1b-e). Additionally, N-cadherin dependent spheroid formation in 67NR mouse breast cancer cells expressing exogenous N-cadherin was blocked by the N-cadherin antagonists (Fig. 1f, g), while there was no effect on N-cadherin negative cells (Supplementary Fig. S1a, b).

**Fig 1.**
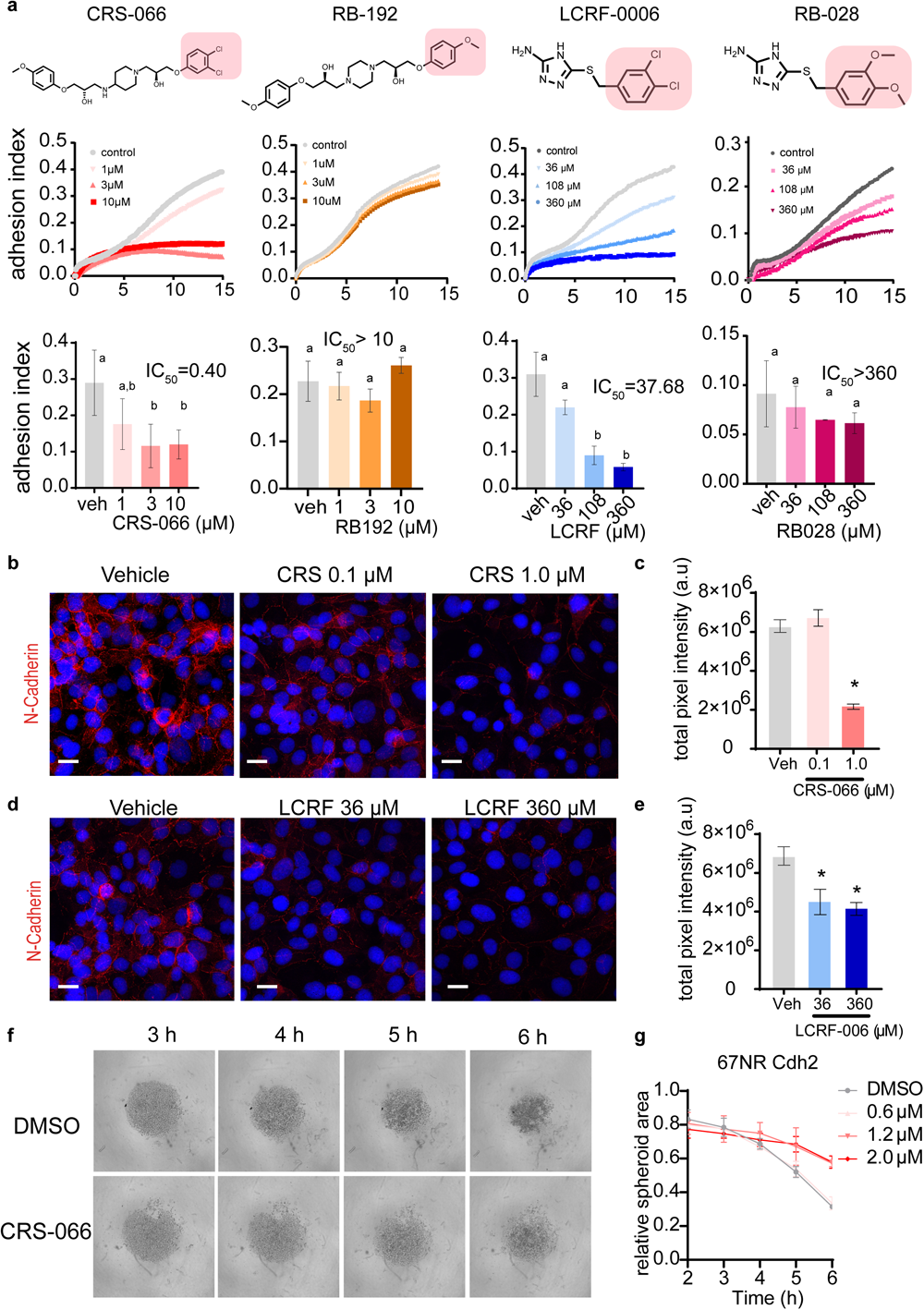
N-cadherin controls adhesive capacity of ovulating COCs. **(a)** top panel: Chemical structures of N-cadherin antagonists CRS-066; LCRF-0006 and analogues RB-192 and RB-028. Active side-chain or modified side-chain depicted by highlighted box. **middle panel:** Average cell adhesion index values of pre-ovulatory COCs (11h post-hCG) interacting with a fibronectin substrate in presence of vehicle or N-Cadherin antagonists CRS-066, LCRF-0006, and sidechain modified analogues RB-192 and RB-028 or vehicle at respective doses. Cell indices were determined using the xCELLigence Impedance system over 15h (n=3 independent experiments with 2 technical replicates per treatment). **bottom panel:** Mean ± SD adhesion index values at 6 h and IC_50_ values of respective drugs calculated from dose-response curve and compared using one-way ANOVA. (**b** and **d)** Representative confocal images of N-cadherin adherens junctions on SK-OV-3 cells treated with N-cadherin antagonists CRS-066 (0.1-1.0 μM), LCRF-0006 (36-360 μM) or vehicle for 24h. (N=3). Scale bar: 40μM.(**c** and **e**) Quantification of N-cadherin adherens junctions. Mean of total pixel intensity in red channel, +/- SEM of at least 50 cells. N=3 independent experiments. Statistical analyses with two-tailed unpaired t-test. * denotes <0.01.(**f**) Bright-field images of spheroid formation in 67NR mouse mammary cell line expressing ectopic Cdh2 were treated with CRS-066 or vehicle at respective time-points. Cells were seeded at 2000 cells per well, and formation of spheroids was assessed by imaging every hour for 6h. (**g**) Mean +/- SEM of spheroid area in 67NR-Cdh2 cells treated with either vehicle or increasing doses of CRS-066 (0-2 μM) over 6h.

Together, these data reveal that N-cadherin plays a hitherto unappreciated role in periovulatory COCs and that these new N-cadherin antagonists can disrupt adhesion of periovulatory COCs to substratum (i.e. fibronectin) as well as blocking cell-cell adhesion in other non-ovarian, N-cadherin expressing cells.

### Adherens complexes between cumulus cells and oocytes are disrupted by N-cadherin inhibition

Expression of *Cdh2*, encoding mouse N-cadherin was equivalently high in granulosa cells and COCs with no significant regulation by hormones inducing follicle growth (44 h eCG), or ovulation, with the exception of a transient 4-fold reduction in *Cdh2* at the time of ovulation (12 h post hCG) specifically in COCs (Fig. 2a). Expression of *Ctnnb1* encoding the adherens junction cytoskeletal adaptor protein β-catenin was high in granulosa and COCs and maintained throughout folliculogenesis and ovulation (Fig. 2b). *Cdh1* encoding mouse E-cadherin was 5-fold more abundant in COCs than granulosa cells with a significant 5-fold reduction in mRNA expression seen 8 h and 12 h after ovulation was induced by hCG (Fig. 2c).

**Fig 2.**
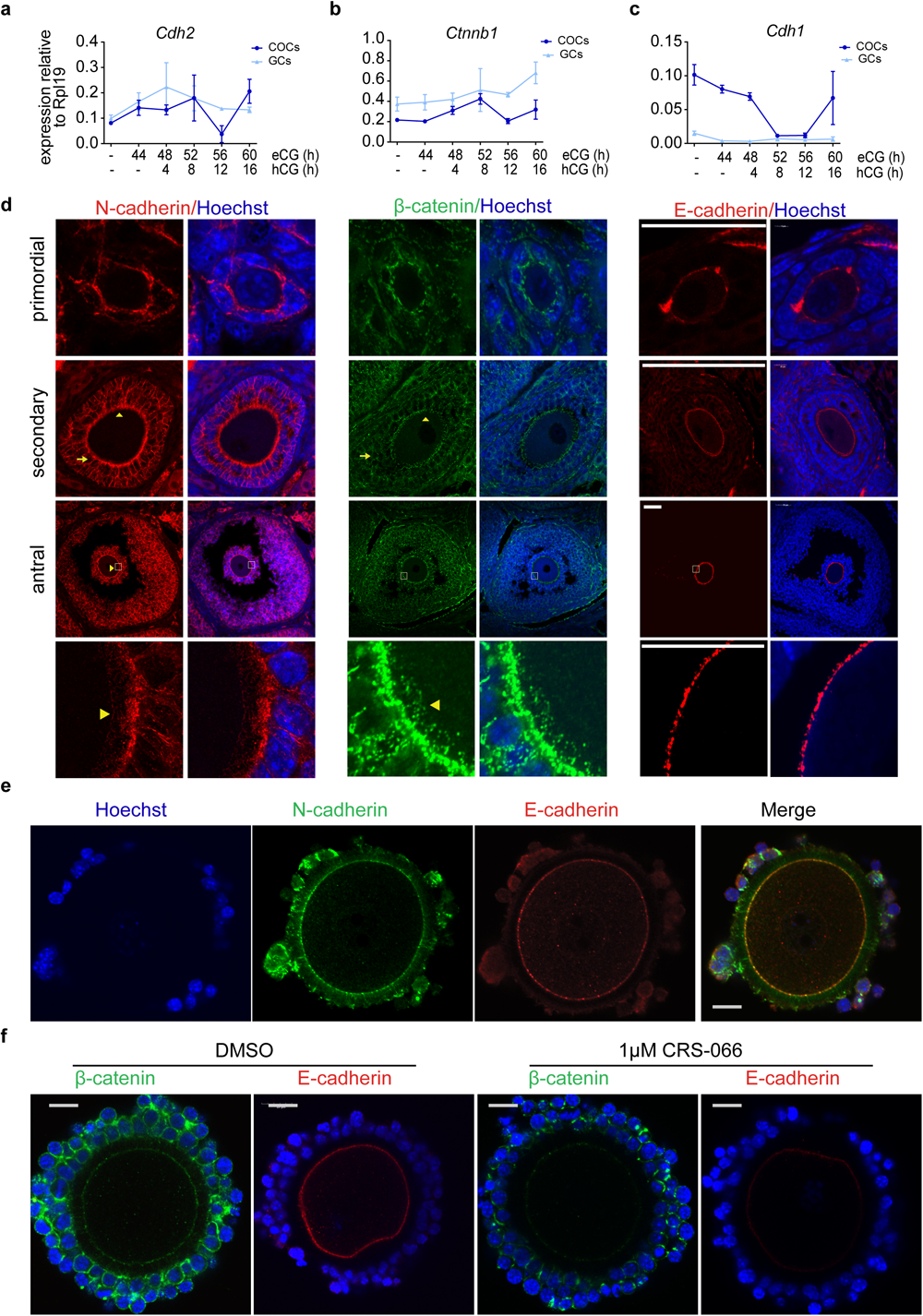
N-cadherin maintains oocyte-cumulus cell interaction. (**a**-**c**) Time-course of Cdh2, ctnnb1 and Cdh1 mRNA expression in isolated granulosa cell (GCs) or Cumulus oocyte complexes (COCs) from mouse ovaries at indicated time-points after eCG and hCG stimulation of folliculogenesis and ovulation (N=3 animals per time point). Cdh2 and Ctnnb1 levels are high in GC and COC throughout folliculogenesis, with a transient drop in Cdh2 level 12 h after ovulation stimulus, while E-Cadherin was high in COCs and significantly reduced by ovulation stimulus. The levels shown of the indicated mRNAs were determined by TaqMan qPCR normalised to Rpl19. (**d**) Immunofluorescent staining of N-Cadherin, βcatenin and E-cadherin throughout ovarian folliculogenesis. Confocal images of mouse ovarian sections obtained from eCG primed mice and stained using anti N-Cadherin (left panel), anti-β-catenin (middle panel) and E-cadherin (right panel). DNA is counterstained with Hoechst. Arrows indicate presence of N-Cadherin and β-catenin at granulosa-granulosa cell junctions in secondary and antral follicle stages. Arrowheads indicate presence of N-cadherin and β-catenin at oocyte-cumulus interface. High magnification images show transzonal projections extending from cumulus cells and anchored to oocyte membrane Scale bar: 50 µM. (**e**) Whole-mount immunofluorescent staining showing co-localization of N-cadherin (green) and E-cadherin (red) at the oocyte plasma membrane in mouse COC from antral follicles of eCG primed mice. N-cadherin is also evident on cumulus cell surfaces and transzonal projections. Cumulus cell and oocyte nuclear DNA is counterstained with Hoechst. Scale bar: 50 µM. (**f**) Wholemount immunostaining shows loss of β-catenin and E-cadherin at the oocyte plasma membrane after treatment with CRS-066. COCs obtained from antral follicles of eCG primed mice and treated with CRS-066 or vehicle for 4h. COCs were fixed and stained with anti-E-cadherin (green) and anti-β-catenin (red). DNA was counterstained with Hoechst. Scale bar: 50 µM.

N-cadherin and β-catenin proteins were abundant on granulosa and cumulus cell surfaces at intercellular junctions (Fig. 2d indicated by arrows) from primordial follicle stage and throughout folliculogenesis and ovulation. In contrast, both theca interna and externa had no detectable N-cadherin. N-cadherin and β-catenin were also abundant along cumulus cell TZPs extending across the zona pellucida and contacting the oocyte membrane (arrowheads Fig. 2d). Interestingly, N-cadherin and E-cadherin were shown to co-localise at the oocyte plasma membrane by co-staining in wholemount COCs (Fig. 2e).

The assembly of functional adherens complexes at granulosa cell interfaces was confirmed using proximity ligation assay (PLA) which showed that N-cadherin and β- catenin are present in the same protein complexes at points of intercellular contact in cultured granulosa cells (Supplementary Fig. S2a). Likewise, in whole-mount COCs the cumulus cells showed numerous foci of interacting N-cadherin and β-catenin at cell-cell contacts between cumulus cells (Supplementary Fig. S2b). Notably, we also found PLA signals for N-cadherin and β-catenin interaction at the oocyte plasma membrane indicating adherens complex formation containing N-cadherin and β- catenin at the cumulus cell-oocyte interfaces (Supplementary Fig. S2c). Since the oocyte contains only E-cadherin, this suggests that heterotypical adherens complexes of N-cadherin and E-cadherin form at the cumulus cell-oocyte interface. Furthermore, we found that cumulus cell N-cadherin is required to stabilise oocyte membrane E-cadherin and β-catenin by incubating COCs with 1μM CRS-066 or DMSO control for 4h. Wholemount immunostaining showed β-catenin and E-cadherin localization at the oocyte membrane in controls, while treatment with CRS-066 caused decreased β- catenin on cumulus cell surface, as well as on the oocyte membrane and a concomitant loss of E-cadherin on oocyte membrane (Fig. 2f).

### N-cadherin antagonists disrupt cumulus expansion and oocyte meiotic maturation

To determine whether the loss of oocyte membrane β-catenin and E-cadherin upon N-cadherin inhibitor treatment disrupts the bi-directional communication between cumulus cells and the oocyte we performed *in vitro* maturation (IVM) in the presence of the N-cadherin antagonists or vehicle controls and assessed COC expansion and oocyte meiotic maturation.

Increasing concentrations of LCRF-0006 (36 – 360 μM) dose dependently reduced cumulus expansion index at the end of 12 h IVM (Fig. 3a, b). This inhibition of COC expansion after treatment with LCRF-0006 was not due to impaired response to the FSH and EGF stimulus or activation of critical genes. Expression of hyaluronan synthase (*Has2*) which produces the key cumulus matrix component hyaluronan, was in fact dose-dependently increased by LCRF-0006 (Fig. 3c). Expression of known mediators of COC expansion amphiregulin (*Areg*) and the prostaglandin synthase enzyme (*Ptgs2*), [32, 33] were either increased or unaffected by LCRF-0006 treatment (Fig. 3c). mRNA for connective tissue growth factor (*Ctgf*), a canonical YAP1 target, was significantly reduced by the two highest doses of LCRF-0006.

**Fig 3.**
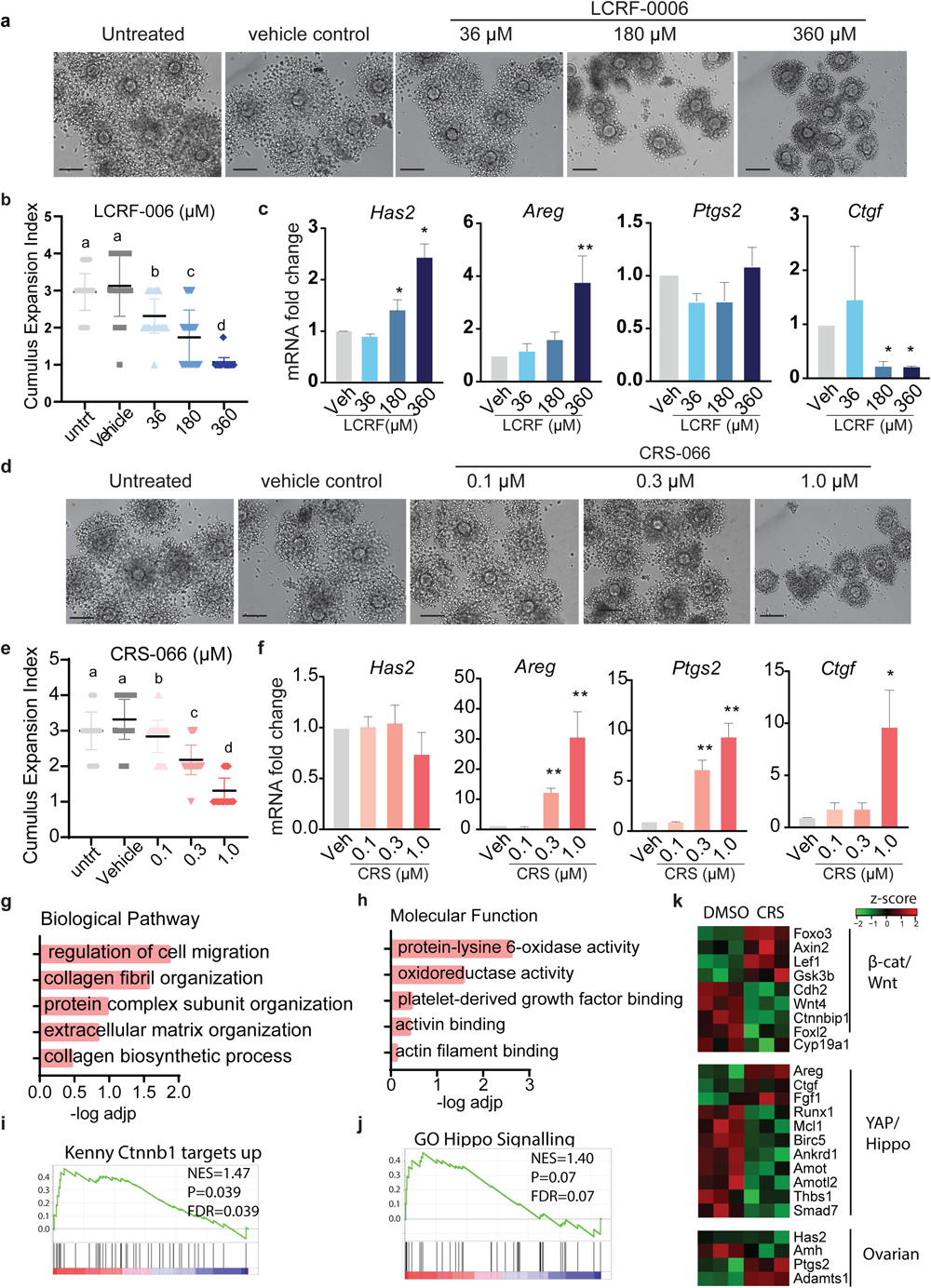
N-cadherin antagonists block cumulus expansion in mouse COCs. COCs from eCG primed mice were treated with LCRF-0006 (36-360 uM) or CRS-066 (0,1-1 uM) during *in vitro* maturation (EGF and FSH stimulated) and cumulus expansion was assessed after 12 h or gene expression assessed after 10 h IVM. (**a** and **d**) Representative bright-field images of COCs after 12 h IVM treated with LCRF-0006 or CRS-066. Scale bar: 10µm. (**b** and **e**) Mean ±SEM of Cumulus expansion indices from a and d n= >20 COCs per experiment. N=4 independent experiments, * = P<0.05, ** = P<0.01. (**c** and **f**) Effect of N-cadherin antagonist treatment during IVM (10 h) on the expression of key genes involved in COC expansion during IVM. Mean± SEM. N=3 independent experiments. Statistical testing with one-way ANOVA * P<0.05, **P<0.01. (**g** and **h**) Gene ontology enrichment of biological pathways and molecular functions of significantly differentially down-regulated genes identified in RNA-Seq analysis of COCs after CRS-066 (0.3 μM) treatment compared to vehicle treatment.. All data are presented as the ratio of CRS-066 over vehicle (N=3). (**I** and **j**) Gene set enrichment analysis plot (GSEA) demonstrating the upregulation of Ctnnb1 and Hippo signalling pathways in CRS-066 treated COCs versus vehicle treated COCs. Net enrichment score (NES) values are shown. N=3 independent biological replicates. (**k**) Heat map representing the relative expression profiles of transcripts involved in Wnt\β- catenin, Hippo\YAP and ovarian signalling axes.

The other N-cadherin antagonist, CRS-066, had a similar effect, leading to a dose-dependent reduction in cumulus expansion during IVM, from 0.1 – 1.0 μM (Fig. 3d, e). Again, the response to IVM stimulus and induction of critical COC expansion genes was not prevented. The expression of *Areg, Ptgs2,* and *Ctgf* were significantly increased by CRS-066 while *Has2* remained unaltered (Fig. 3f).

RNA-Seq was performed to determine the effect of the N-cadherin antagonists CRS-066 n on global gene expression during maturation of COCs. Principal component analysis (PCA) showed separate clusters of gene expression in CRS-066 treated and vehicle controls, indicating a consistent effect on cell signalling by the antagonist (Supplementary Fig. S3a). Differential expression analysis confirmed 168 significant differentially expressed genes (DEGs) (adjusted p-value <1×10^-6^, log2 fold change ≥ ±0.5), with 37 genes downregulated and 131 upregulated in CRS-066 treated COCs (Supplementary Fig. S3b). Gene ontology analyses of the DEGs down-regulated by CRS-066 treatment showed that regulation of cell migration and ECM reorganisation were the most over-represented biological pathways (Fig. 3g). Protein-lysine 6-oxidase activity, an enzyme responsible for cross-linking of collagens, was the most over-represented molecular function (Fig. 3h). Gene set enrichment analysis (GSEA) identified the β-catenin and Hippo signalling pathways canonical downstream signalling pathways of N-cadherin were significantly enriched in CRS-066 treated COC DEGs (Fig.3i, j). In agreement with the qPCR analysis, the RNA-seq results showed treatment with CRS-066 increased the expression of Hippo-pathway responsive genes including *Areg* and *Ctgf* (Fig. 3k). In contrast, the expression of critical granulosa cell-specific and β-catenin responsive genes *Foxl2* and *Cyp19a1*, were significantly downregulated in COCs with CRS-066 treatment.

To confirm the notion that YAP1 is required for COC maturation we assessed the effect of YAP1 inhibition on cumulus expansion during IVM. Treatment with YAP inhibitor, Verteporfin (3 μM), significantly reduced COC expansion with concomitant decrease in the canonical YAP1 target *Ctgf* as well as *Areg*, and *Ptgs2,* while *Has2* expression remained unaffected. These gene expression changes are all consistent with increased YAP1 activity in CRS-066 treated COCs (Supplementary Fig. S3c).

Our findings that N-cadherin is present in intercellular junctions between cumulus cells and oocytes, and that N-cadherin antagonists alter cumulus cell response during IVM, led us to investigate the impact of the antagonists on oocyte meiotic maturation. Both CRS-066 and LCRF-0006 treatment during IVM dose dependently reduced the proportion of oocytes undergoing germinal vesicle breakdown (GVBD) and the proportion reaching MII stage, as determined by the presence of an extruded polar body after 12 h IVM (Fig. 4). The stage of meiotic arrest was determined by visualizing meiotic spindles and cortical actin in oocytes by labelling β-tubulin, actin staining (phalloidin), and chromosomes (Hoechst). In untreated and vehicle treated controls >90% of oocytes reached MII stage, as demonstrated by successful assembly of the metaphase II spindle and extrusion of the sister chromatid into polar bodies. Increasing doses of CRS-066 caused more oocyte arrest at GV-intact or MI stage (Fig. 4a-c). Specifically, 0.3 μM CRS-066 significantly reduced polar body extrusion (PBE) to 57%, with over 30% oocytes arrested in MI, while 1.0 μM CRS-066 resulted in more than 50% oocytes arresting at GV and the remainder after GVBD. Oocytes exhibiting MI spindle formation were rarely observed. LCRF-0006 treatment also caused oocyte arrest in early meiosis, with 20% arrested in GV stage at the lowest 36 μM dose and almost all oocytes arresting with intact GVs at the highest (360 μM) dose (Fig.4d-f).

**Fig 4.**
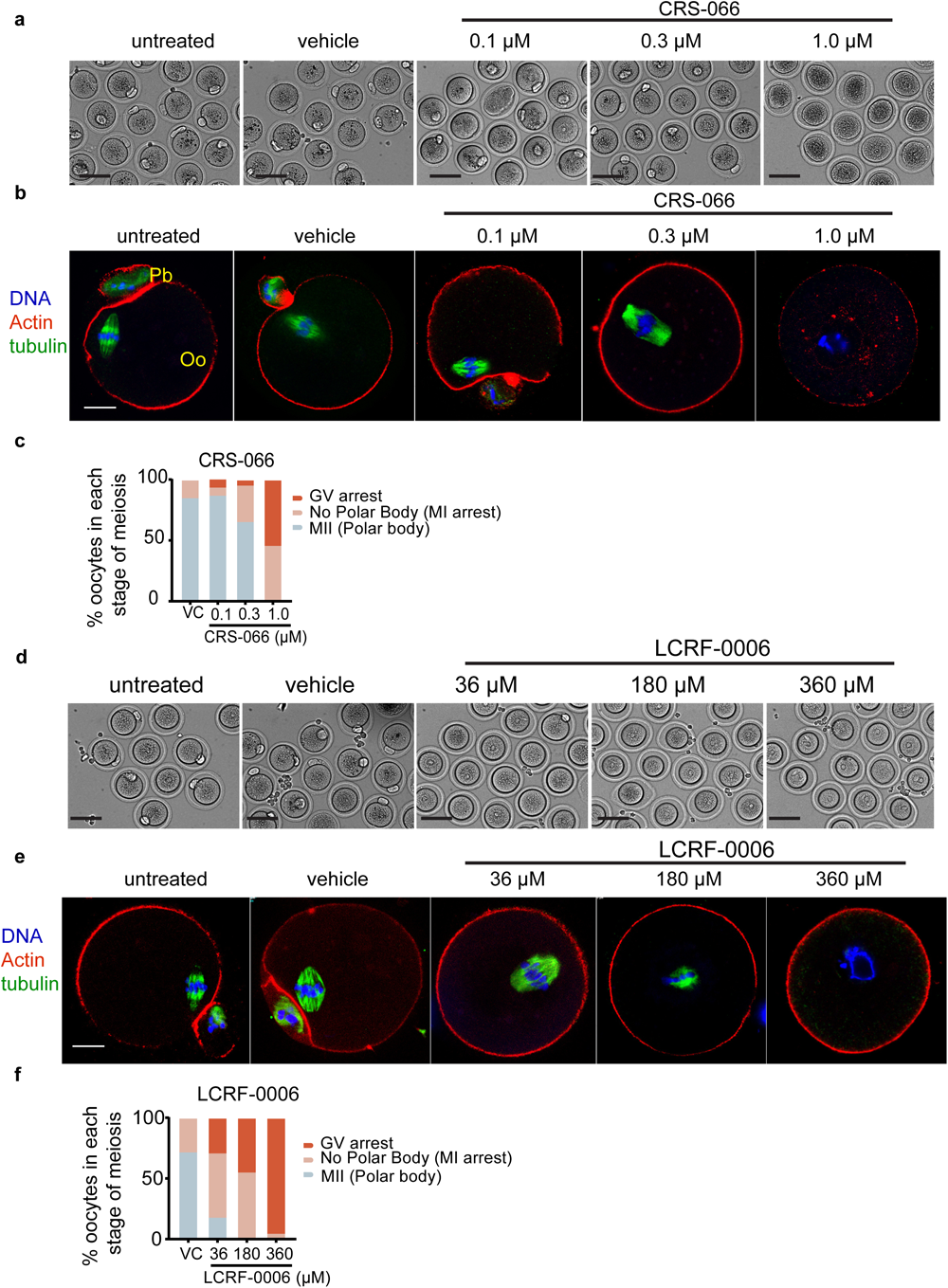
N-cadherin antagonists block meiotic maturation in mouse COCs. COCs from eCG primed mice were treated with LCRF-0006 (36-360 uM) or CRS-066 (0,1-1 uM) during *in vitro* maturation (12 h EGF and FSH stimulated), then oocytes were denuded and meiotic stage was assessed by labelling actin (phalloidin, red), spindles (tubulin IF, green) and DNA (Hoechst, blue). n= >20 COCs per experiment; N=4 independent experiments. (**a** and **d**) Representative bright field images of denuded mouse COCs showing oocyte and polar body morphology after vehicle, CRS-066 or LCRF-0006 treatment. Scale bar: 5µM. (**b** and **e**) Representative confocal fluorescent images of denuded mouse oocytes after IVM treated with LCRF-0006 or CRS-066 as indicated, showing polar bodies, spindle structure or germinal vesicle morphology typical in each treatment condition. Scale bar: 20µM. Pb indicates polar body formation; Oo indicates Oocyte. (**c** and **f**) Percent of oocytes at each stage of meiotic progression from n>20 oocytes per experiment in 4 independent experiments. MI defined by the any evidence of tubulin staining in the first metaphase spindle with no polar body extrusion; MII defined by presence of a polar body and MII metaphase plate. GV defined by oocyte DNA lacking any tubulin staining to indicate MI spindle formation.

### N-cadherin antagonist CRS-066 inhibits ovulation in mice

Given the robust effect of CRS-066 and LCRF-0006 on COC adhesion, expansion and meiotic maturation *in vitro*, we hypothesised that these N-cadherin antagonists may interfere with ovulation *in vivo*. To assess this, stimulated ovulation was performed in prepubertal (3-week old) female mice in conjunction with administration of 2x daily i.p. injections of CRS-066 (50 mg/kg), or LCRF-0006 (100 mg/kg) or vehicle for 4 days (Supplementary Fig. S4a).

CRS-066 treatment significantly compromised the number of oocytes retrieved from oviducts after 16 hCG treatment (11.6 ± 7) compared to vehicle treated control mice (26.2 ± 6 ovulations / ovary, p<1×10^-5^, n=6 mice per group) (Fig. 5a). Histological analysis showed the presence of multiple ovulated luteinized structures in vehicle-treated mice, while CRS-066 treated mice had unruptured antral follicles with entrapped oocytes, confirming that ovulation was specifically disrupted by CRS-066 treatment (Fig. 5b). Next, RNA-Seq and differential gene expression analysis in CRS-066 and vehicle-treated whole ovary RNA was performed. Unsupervised hierarchical clustering of RNA-sequencing data identified distinct gene expression profiles in CRS-066 treated compared to control ovaries (Fig. 5c). A total of 330 significant DEGs were identified with (adjusted p-value <1×10^-6^, log2 fold change ≥ ±0.5), with 136 genes upregulated and 194 genes down-regulated (Supplementary Fig. S4b). Gene ontology analyses of genes downregulated by CRS-066 treatment showed that ECM organisation and cell-cell adhesion, migration and motility were the most affected pathways (Fig. 5d). Integrin binding and cadherin binding were the most affected molecular functions with CRS-066 treatment (Fig. 5e). Gene set enrichment analysis (GSEA) showed a profile enriched for both β-catenin target gene set and Hippo signalling pathways (Fig. 5f, g). Of note, CRS-066 repressed the expression of a group of ovarian genes known to be in the β-catenin/Wnt signalling pathway, including *Cyp19a1* (gene encoding aromatase which is involved in estrogen synthesis) that was repressed >2 fold, while *Foxl2*, a forkhead transcription factor specifically expressed in granulosa cells, and Amh, a granulosa specific hormone were upregulated >1.5 fold. Other β-catenin/Wnt signalling pathway genes, including Wnt inhibitory factor (*Wif1*), and Glycogen synthase kinase (*GSK3β*) which phosphorylates β-catenin were also down-regulated after CRS-066 treatment (not shown). CRS-066 treatment also repressed Hippo pathway genes *Areg* and *Ctgf* (Fig. 5h).

**Fig 5.**
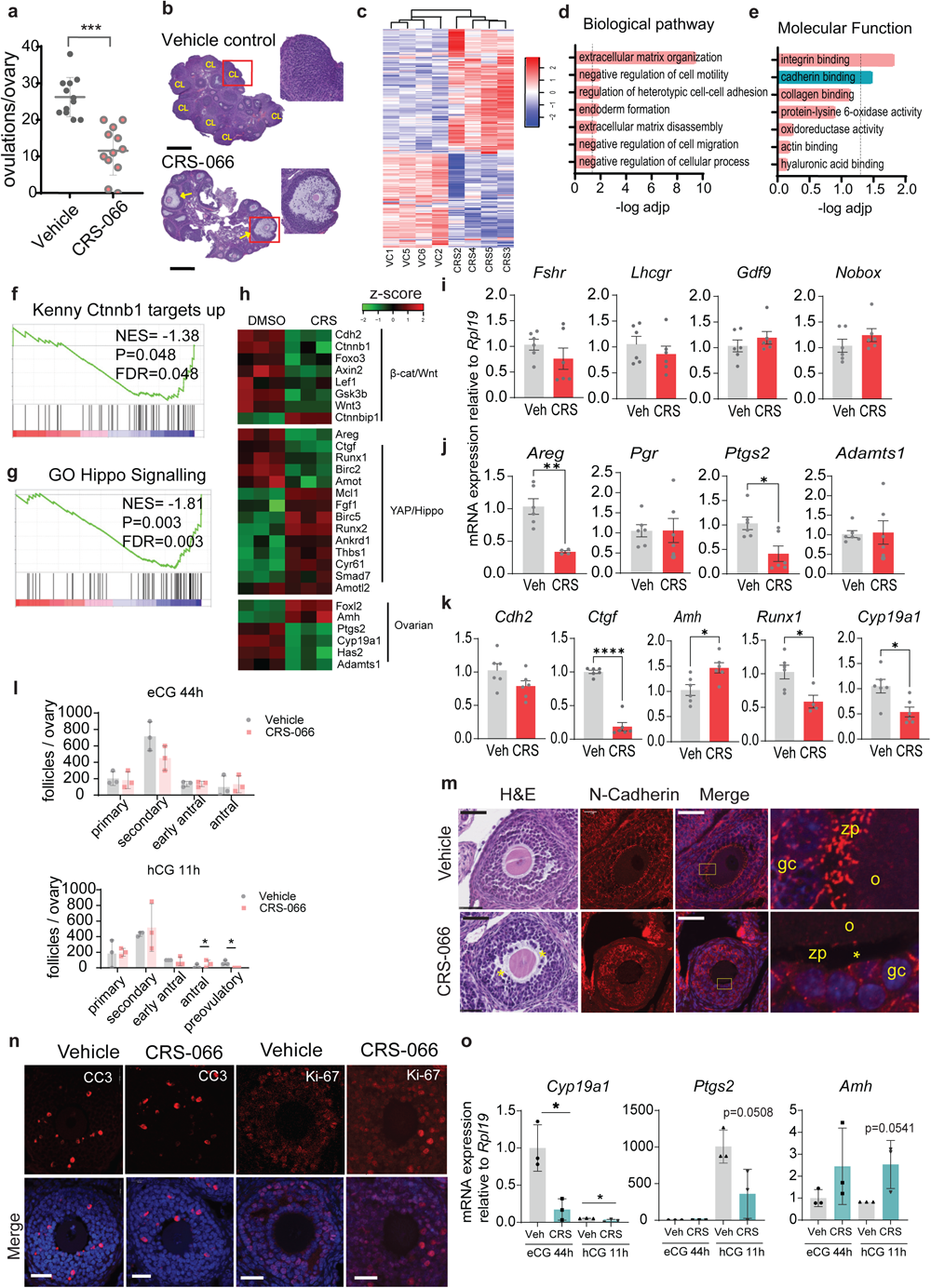
N-Cadherin antagonist CRS-066 blocks ovulation *in vivo*. (**a**) Ovulation rate of 21d old mice treated with CRS-066 (50mg/kg) or vehicle (7.5% DMSO in 0.9% saline). COCs in oviducts counted 16h after hCG injection. Graph represents mean ±SEM from N=6 animals; ***p<0001 (unpaired two-tailed t-test). (**b**) Histology of ovaries by Haematoxylin & Eosin staining. CL indicate corpus leuteum; arrows indicate trapped oocytes in CL. Scale bar: 100µm. (**c**) Hierarchical clustering of RNA-sequencing analysis results shows differentially expressed genes between CRS-066 and vehicle treated mice (N=6 mice per treatment). (**d** and **e**) Gene ontology enrichment of biological pathways (D) and molecular functions (E) of significantly differently down-regulated genes in CRS-066 treated mouse ovaries compared to vehicle treated ovaries. (**f** and **g**) Gene set enrichment analyses plot (GSEA) shows downregulation of Ctnnb1 and Hippo signalling pathways in CRS-066 treated mice ovaries compared to vehicle treated ovaries. Net enrichment score (NES) values are shown. (**h**) Heatmap representing the relative expression profiles of transcripts involved in Wnt\β-catenin, Hippo\YAP and ovarian signalling axes in CRS-066 treated ovaries compared to vehicle. N=3 biological replicates. (**i**-**k**) Relative mRNA expression of key genes involved in gonadotrophin signalling and oocyte function (i), COC expansion and ovulation (j) or folliculogenesis (k) h in ovaries treated with CRS-066 compared to vehicle treatment and determined by qRT-PCR. Bar graph show mean+/- SEM. N=6 ovaries from independent CRS or vehicle treated mice. Statistical testing with Student’s t-test; *P < 0.05; ** P<0.01; **** P <0.00001. (**l**) Follicle counts at primary, secondary, pre-antral, antral and ovulatory stages in ovaries from mice treated with either CRS-006 (50mg/kg) or vehicle control (7.5% DMSO). N=3 mice/treatment/time-point. (**m**) Representative follicle morphology H&E (left) section and N-cadherin immunofluorescence (right) section in mice treated with either CRS-066 or vehicle. H&E and immunoflourescence highlight disorganised granulosa cells organisation. Asterisks indicate loss of transzonal projections between oocyte and cumulus cells. Scale bar: 30µm. (**n**) Representative confocal immunofluorescent images of mouse ovaries stained with anti-cleaved caspase 3 and anti-Ki-67. Scale bar: 30µm. (**o**) Relative mRNA expression of key genes involved in oocyte growth and ovulation in CRS-066 or vehicle treated mice (N=3/ treatment) at either 44h post eCG or 11h post hCG.

Consistent with transcriptomic analysis, qPCR showed that genes involved in granulosa cell hormone responsiveness including the gonadotropin receptor genes (*Fshr*, *Lhcgr*) were unaffected (Fig. 5i), while an elevation in *Amh* and *Foxl2* ((Fig. 5k) expression in whole ovary RNA likely indicates elevated granulosa cells numbers in CRS-066 treated ovaries that fail to ovulate. Expression of oocyte specific genes *Gdf9* and *Nobox* were unaffected by CRS-066 treatment (Fig. 5i). Among genes required for ovulation, *Areg* and *Ptgs2* expression was significantly reduced, while *Pgr* and *Adamts1* were unaffected (Fig. 5j).

The numbers of follicles at each developmental stage were equivalent in CRS-066 and vehicle treated mice. Ovaries from mice treated in the same manner and collected either before hCG treatment (eCG 44 h) or 11 h after hCG (1 h before ovulation) showed equivalent numbers of follicles at each stage of development from primary to antral. Typical periovulatory follicle morphology and number confirmed that the response to hCG was normal (Fig. 5l). Immunostaining of N-cadherin showed a disruption to the normal organisation of N-cadherin at intercellular boundaries in the granulosa layers, loss of N-cadherin in the TZPs and dissociation of cumulus cells from the cumulus – oocyte interface in follicles of mice treated *in vivo* with CRS-066 (Fig. 5m). Proliferation and apoptosis in granulosa cells, assessed by Ki67 and cleaved caspase 3 (CC3) immunostaining were not significantly different between ovaries of vehicle and CRS-066-treated mice (Fig. 5n). The effects of CRS-066 treatment on expression of specific genes in ovaries collected 44h after eCG confirmed a normal response to hormone treatments with expression of Cyp19a1 high in preovulatory (eCG 44 h treated) ovaries, and repressed by hCG stimulation of ovulation, while Ptgs2 expression was low at preovulatory stage, but induced by hCG 11 h. Treatment with CRS-066 at both pre-and periovulatory stages of folliculogenesis caused a significant reduction in Cyp19a1 and Ptgs2, but an increase in Amh expression compared to vehicle controls (Fig. 5o), consistent with the observations in hCG 16 h post-ovulatory ovaries.

Treatment of mice with 100 mg/kg LCRF-0006 for 4 days prior to ovulation resulted in similar numbers of ovulated oocytes retrieved from oviducts (23.9 ± 11.2) compared to vehicle controls (25.6 ± 5.3 ovulations per ovary. P=0.67, n=6 mice per group) after 16 h hCG treatment. Additionally, there were no significant changes in *Cyp19a1*, *Ptgs2* or *Areg* gene expression, indicating this compound was less active *in vivo* as was also the case *in vitro*.

### Granulosa specific N-cadherin knockout inhibits ovulation in mice

To verify that N-cadherin is the specific target of the antagonists that modulate mechanosensitive gene expression and ovulation, we generated granulosa-specific Cdh2 knockout. Breeding crosses of *Cdh2^Fl/+^;Amhr2^Cre/+^* mice producing offspring with the expected genotypes at Mendelian ratios. Female offspring with *Cdh^Fl/+^;Amhr2^Cre^*, versus *Cdh^Fl/FL^;Amhr2^Cre^* (granulosa-specific N-cadherin depletion) were comparatively assessed in pre-ovulatory (eCG 44 h) ovaries. qPCR analysis confirmed significant 4-fold reduction in Cdh2 mRNA expression in *Cdh^Fl/FL^;Amhr2^Cre^*ovaries (Fig 6a). Likewise, immunofluorescence of ovarian sections showed dramatically reduced N-cadherin protein in ∼80-90% of granulosa cells in all follicles. A mosaic pattern was observed with 10-20% of granulosa cells in antral follicles showing approximately normal levels of N-cadherin immunostaining (Fig 6b-e), consistent with other reports of incomplete penetrance of the Amhr2-cre transgene expression in granulosa cells [34, 35]. The N-cadherin depleted areas of follicles showed multiple regions of detached cells suggesting a loss of intercellular adhesion. The expression of putative N-cadherin mechanoresponsive genes Areg and Ptgs2 were significantly (P<0.05) reduced 2-3-fold respectively in the granulosa-specific Cdh2 mutants, and Cyp19a1 showed trending (P=0.08, 2.5-fold) reduced expression (Fig 6a), similar to the reduced Cyp19a1 expression reported in *Ctnnb1^Fl/FL^;Amhr2^Cre^*mice [35]. Ovulation examined by collecting and counting COCs in oviducts of female *Cdh^Fl/FL^;Amhr2^Cre^*mice after eCG+hCG 14h stimulation was significantly (P<0.01, 3-fold) reduced (Fig 6f) and histological assessment identified multiple large unruptured follicles in the *Cdh^Fl/FL^;Amhr2^Cre^* granulosa specific Cdh2 mutant ovaries (Fig 6g). Together these results support the conclusion that N-cadherin is important for cell-cell adhesion and mecahnosignalling within granulosa cell populations, and for successful ovulation of the mature preovulatory follicles.

**Fig 6.**
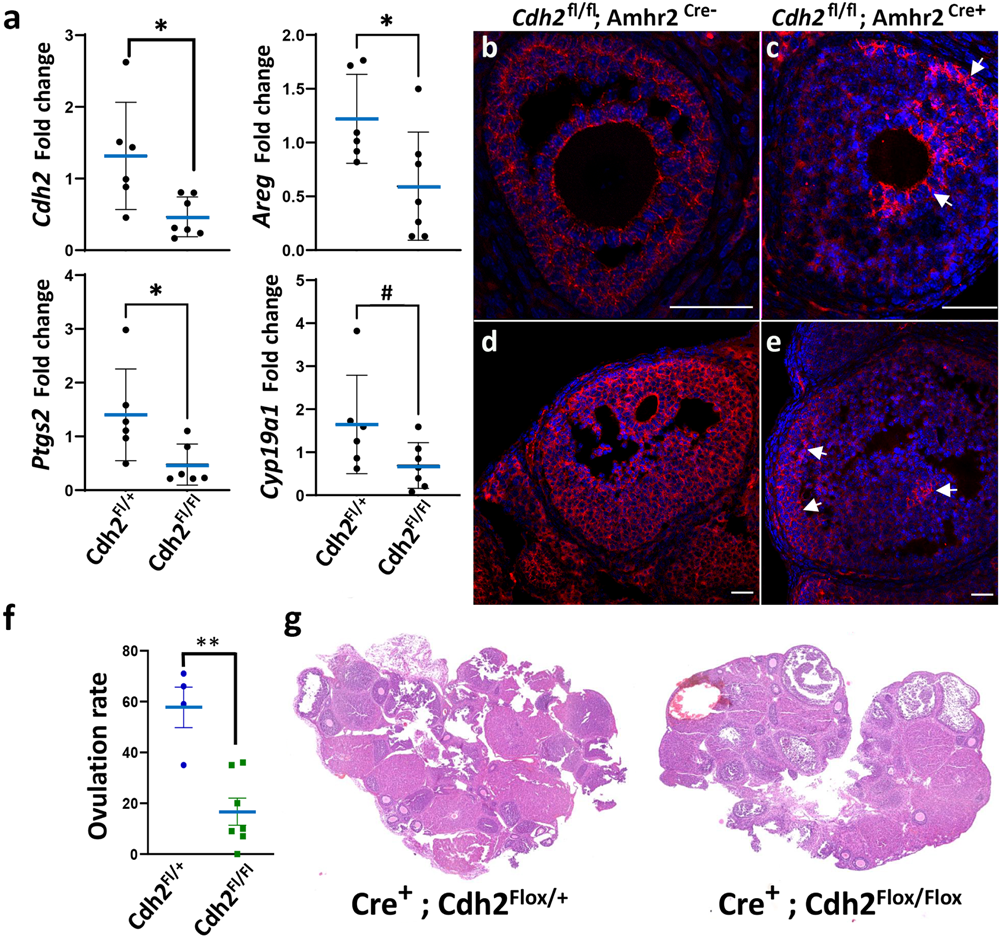
Granulosa conditional Cdh2 null mutation disrupts ovarian gene expression and blocks ovulation. **(a)** qPCR analysis of relative mRNA expression of Cdh2, Areg, Ptgs2 and Cyp19a1 in ovaries of control (*Cdh2^Fl/+^ ; Amhr2^Cre^*) and granulosa-specific Cdh2 null mutants (*Cdh2^Fl/Fl^ ; Amhr2^Cre^*), n=6 individual animals, *p<0.05. (**b-e**) Immunofluorescent analysis of N-cadherin protein in ovaries of control (*Cdh2^Fl/+^ ; Amhr2^Cre^*) and granulosa-specific Cdh2 null mutants (*Cdh2^Fl/Fl^ ; Amhr2^Cre^*), showing mosaic depletion of N-cadherin in granulosa cells of mutant follicles. Arrows indicate mosaic regions with persistent N-cadherin. (**f**) Ovulation rate of 21d old mice with indicated control or granulosa-specific mutant genotypes. COCs in oviducts counted 16h after hCG injection. Graph represents mean ±SEM from N=4 and 7 animals respectively; **p<0.01 (unpaired two-tailed t-test). (**g**) Histology of ovaries by Haematoxylin & Eosin staining. Scale bar: 100µm.

## DISCUSSION

Our study shows, for the first time, that N-cadherin plays a central role during late follicular morphogenesis. Specifically, N-cadherin antagonists, or Cdh2 gene mutation disrupt intercellular contacts between granulosa cells, as well as oocyte-cumulus cell contacts leading to impaired COC expansion, oocyte maturation and ovulation. Likewise, even incomplete disruption of the Cdh2 gene expression led to a similar disruption to mechanoresponsive gene expression and ovulation. Broad transcriptional changes indicate cross-talk between N-cadherin with β-catenin and Hippo/YAP pathways mediating a mechanical signal that integrates with the hormone actions to regulate genes required for oocyte maturation and ovulation.

There is evidence that the ovarian follicle is a mechanosensitive organ. Either excess or inadequate tissue rigidity profoundly affects folliculogenesis and oocyte maturation. This is evident in Polycystic Ovarian Syndrome (PCOS), a prevalent infertile condition where excess rigid ECM prevents follicle and oocyte growth progression [36]. It is also shown *in vitro*, where isolated follicles in different bioengineered support matrices exhibit reduced follicle growth and viability in soft or rigid environments [37]. Our findings suggest that N-cadherin is a key mechanosensor and transducer of growth and differentiation of granulosa cells, cumulus cells and oocytes. Consistent with this, dysregulation of β-catenin and Hippo/Yap1 has been shown in conditions of disrupted follicle growth including PCOS [22], as well as Primary Ovarian Insufficiency (POI) and fibrosis [38, 39]. Further studies are required to determine the interplay between cadherins and integrins in the cells of ovarian follicles, as both cell surface receptor groups interact with the actin-based microfilaments and regulate similar biological processes.

Direct physical contacts between cumulus cells and the oocyte are important for transport of cAMP and cGMP via gap junctions to maintain oocyte meiotic arrest. Both *in vivo* and *in vitro* N-cadherin antagonists caused disruption of transzonal projections and cumulus-oocyte cellular contacts in COCs. Disruption of these contacts was recently shown to be a normal event in response to induced COC maturation [12]. The effects of N-cadherin antagonists provide the first evidence confirming that N-cadherin stabilises these unique cell-cell contacts and that premature disruption of the cumulus cell-oocyte contacts is highly detrimental to follicle growth and oocyte maturation. Our observation that two N-cadherin antagonists severely impaired COC expansion and meiotic maturation in IVM while not depleting expression of cumulus matrix genes such as *Has2* suggests that N-cadherin mediates signalling between oocyte and cumulus cells, transducing essential signals necessary for these pre-ovulatory events. This is in line with N-cadherin being a known mechanotransducer and being abundant at cell junctions between cumulus cells and in transzonal projections at cumulus-oocyte contacts. The disruption of cumulus cell arrangement around oocytes in ovaries of N-cadherin antagonist treated mice also demonstrates the necessity for N-cadherin to stabilise cell-cell contact and maintain the tissue architecture critical for bidirectional cell communication. The possibility of heterotypic interaction of N-and E-cadherin at the cumulus cell-oocyte interface suggests a unique type of intercellular junction in the context of oocytes and cumulus cells with interactomes that are also likely to be specialised. The molecular composition of these adherens complexes, and the mechanism behind cadherin driven intracellular signalling in the COC clearly warrants further investigation.

β-catenin transduces Wnt signalling, and in ovarian cells it is fundamental to specification of the female somatic cell lineage [19] establishing female sex determination and follicle formation (reviewed in [27]). Our studies are the first to demonstrate that N-cadherin is important for functional gene regulation in the follicle by regulating the β-catenin and Hippo/Yap1 pathways. Reduced expression of β- catenin regulated granulosa cell specific genes, including *Ptgs2* and *Cyp19a1* [20, 35, 40, 41], and changes in the transcriptome profile indicate the loss of β-catenin activity caused by the N-cadherin antagonist. This is consistent with N-cadherin’s reported role stabilising β-catenin at tensile cell junctions and facilitating nuclear translocation in response to Wnt signals [42, 43]. Thus, our results demonstrate that tissue structure and mechanical force contribute to hormone responsiveness and the steroidogenic phenotype of granulosa cells. We did not see evidence of upregulated testis specific genes indicating that neither N-cadherin antagonist treatment, nor gene depletion was not sufficient to reverse granulosa cell specification as seen with complete β-catenin ablation [44]. However, our results do indicate that N-cadherin stabilises granulosa and cumulus cell phenotypes through β-catenin signalling.

N-cadherin antagonists also reduced expression of Yap1 target genes with key functions in folliculogenesis (*Ctgf*, *Areg*) [43], indicating a loss of Yap1 function *in vivo*. These findings suggest that an ovarian Yap1 mechanosensing pathway, also mediated through N-cadherin, is attenuated in the antagonist treated mice. Trans-cellular interaction of cadherins has been shown to sequester Yap1 in the cytoplasm [46–48], and peptide inhibitors of N-cadherin can block nuclear translocation of Yap1 in response to tissue rigidity in mesenchymal stem cells [49]. Thus, our results show Yap1 signalling in granulosa cells is controlled by N-cadherin and regulates the differentiation status of these cells and their response to hormone stimuli. This conclusion is also supported by the morphological change in follicles of Wnt4 deficient mice [13], and in granulosa-specific Yap1 knockout mice [45]. Together, these findings suggest that mechanotransduction signals through N-cadherin mediate the reported high Yap1 activity in granulosa cells needed to sustain normal folliculogenesis.

During *in vitro* maturation of COCs, N-cadherin antagonists blocked COC expansion, and oocyte meiosis. CRS-066 paradoxically increased *Areg* and *Ptgs2*, which are β- catenin repressed genes in COCs [20]. Their induction is consistent with compromised β-catenin function after N-cadherin inhibition *in vitro* as was seen *in vivo*. The induction of *Areg*, *Ptgs2* and the canonical Yap1 target, *Ctgf* all suggest activation of Yap1 by N-cadherin antagonist in IVM. Supporting this, the Yap1 antagonist verteporfin caused dose-dependent repression of *Ctgf*, *Areg* and *Ptgs2* indicating Yap1 indeed mediates induction of these critical COC maturation genes. This is consistent with reports that show that close contact with oocytes suppresses Yap1 activation in cumulus cells, and that cumulus expansion elicits a mechanical signal that releases this inhibition leading to expression COC maturation genes [21]. Thus, N-cadherin antagonist-mediated inhibition of mechanotransduction disrupted the normal regulation of Yap1. Ctgf mediates size regulation of many organs and is a known ovarian mitogen that directly regulates follicle development and ovulation by upregulating activin-dependent LOX mediated extracellular remodelling, [50]. N-cadherin antagonism during IVM and *in vivo* consistently showed that Lysyl oxidase, activin binding and ECM remodelling were amongst the most affected molecular functions, further supporting our proposed model of N-cadherin acting through Hippo/Yap1/Ctgf to regulate COC expansion, meiotic maturation and ovulation. The inverse effect of LCRF-0006 on *Ctgf* expression in IVM suggests the two antagonists differ not only in their potency but have different effects on N-cadherin signalling responses. Further investigation is required to determine whether this observation is explained by the targeting of different binding pockets of N-cadherin by LCRF-0006 and CRS-066. Since Yap1 and β-catenin have divergent effects on *Ptgs2* and *Areg and* are both influenced by N-cadherin antagonists, the activation/inactivation state of these individual pathways is difficult to unravel, however it is clear the net effect mediated through N-cadherin cell contacts plays a key role in mediating follicle growth and differentiation.

In conclusion, our study identified N-cadherin as a mechanosensory regulator important in ovarian granulosa cell differentiation and response to hormone stimuli both *in vivo* and *in vitro*. The action of N-cadherin through β-catenin and Yap1 signal transducers places new emphasis on the role of these pathways as mediators of hormone responses that also incorporates mechanical cues such as the size and physical tension within the ovary, which is influenced by the neighbouring growing follicles as well as ECM composition. Dysregulation of this mechanism has important implications for conditions such as premature ovarian failure, PCOS and fibrosis as well as other forms of unexplained infertility. Additionally, the potential to target this signalling pathway with small molecule inhibitors, to block ovulation is an illustration the principle of non-hormonal ovulation blocking contraceptives.

## METHODS

### Reagents and antibodies

Unless otherwise stated, reagents were purchased from Sigma-Aldrich (St. Louis, MO, USA).

### Study approval

All animal procedures were approved by the University of Adelaide Animal Ethics Committee (Ethics approval IDs M-2018-071 and M-2018-117) and conducted in accordance with the Australian Code of Practice for the Care and Use of Animals for Scientific Purposes.

### Animals and drug treatments

Hybrid F1 (CBA female/C57 male) prepubertal female mice (21 day old, weighting 10-12 g) were purchased from Laboratory Animal Services (University of Adelaide) and were maintained on a 12 h light and 12 h dark cycle, with rodent chow and water provided *ad libitum*. Mice were injected intraperitoneally (i.p.) with equine chorionic gonadotropin at 5 IU/0.1 ml saline (0.9%) per 12 g of bodyweight, followed 44 h later by i.p. injection of human chorionic gonadotropin at 5 IU/0.1 ml saline per 12 g of bodyweight. The N-cadherin antagonists CRS-066 and LCRF-0006 were a kind gift from Zonula Inc., Kirkland, Quebec, Canada. CRS-066 (compound-15 di-succinate salt) was injected i.p. at 50mg/kg in 0.05 ml saline and LCRF-0006 was injected i.p at 100 mg/kg of in 0.05 ml saline containing 40% 2-hydroxyproprol-β-cyclodextrin twice daily for four consecutive days starting the day prior to treatment with eCG.

### Conditional granulosa Cdh2-null mice

Cdh2 Floxed mice (B6.129S6(SJL)-*Cdh2^tm1Glr/J^*) with exons 1, containing the translational start site of the Cdh2 gene flanked by two loxP sites [51, 52] were obtained from Jackson Laboratories. Mice were genotyped by performing PCR using the following primers: Cdh2F 5’-CCAAAGCTGAGTGTGACTTG-3’ and Cdh2R 5’- TACAAGTTTGGGTGACAAGC -3’ To generate animals with Cdh2 gene ablation Cdh2 floxed mice were crossed with Amhr2-Cre knock in mice (B6;129S7- *Amhr2^tm3(cre)Bhr^*[52] genotyped by PCR using the following primers: Amhr2-Cre-F 5′- CCTGGAAATGCTTCTGTCCG -3′ and Amhr2-R 5′- CAGGGTGTTATAAGCAATCCC-3′.

### Cell line and growth conditions

The human ovarian cancer cell lines SK-OV-3 and 67NR were cultured in Dulbecco’s modified Eagle’s media supplemented with 10% (v/v) FCS, 2 mM ʟ-glutamine, 100 U ml^−1^ penicillin and 100 µg ml^−1^ streptomycin at 37 °C in an atmosphere of 5% CO_2_ in air.

67NR cells were transduced with C-terminal EGFP tagged murine N-cadherin lentiviral construct. The N-cadherin-EGFP construct was cloned into the pCDH-CMV-CMV-MCS-EF1a-Puro lentivirus vector. Cells were transduced with 5 x 10^4^ IU/ml and selected by flow cytometry using EGFP fluorescence to isolate N-cadherin expressing cells.

### xCELLigence real-time cell adhesion assay

Assay of the induced COC adhesion to fibronectin was performed using the xCELLligence system (ACEA Biosciences) in E-plate 16 according to the manufacturer’s instruction as previously described with little modification [26]. Briefly, The E16 xCELLigence plates were prepared by coating with 50μl of mouse recombinant fibronectin (5ug/ml) diluted in αMEM medium before incubation for a minimum of 1 h at 37°C/5% CO_2_. Wells were washed twice with 100 µL/well of αMEM medium. Drugs were added to each well at required concentrations in 50µl bicarbonate buffered αMEM/1% FCS medium. Plates were inserted into the xCELLigence station, and the base-line impedance was measured. Freshly isolated COCs from CBA-F1 mice at 11 h post-hCG, just prior to ovulation, were seeded into each well in 50µl of bicarb buffered αMEM and electiral impedance was measured every 5 mins until 15 hours after seeding. Each treatment was run in duplicate and each plate contained untreated COC control wells. Results are presented as adhesion index over time as well as the concentration eliciting 50% inhibition of adhesion response for N=3 independent biological replicates.

### Spheroid formation assay

Spheroid assay was conducted in 96 well U bottomed ultra-low adhesion Nunclon Sphera plates (Thermo Scientific). Sub-confluent N-cadherin-EGFP 67NR cells were dissociated with TrypLE, washed with warm PBS and counted with the Countess cell counter (Invitrogen). Cells were seeded at 2000 cells per well and centrifuged at 300g for 3 minutes before being placed in culture and imaged every 60 minutes from 2-6 h after seeding on the Incucyte live cell imager (Essen Bioscience) using the spheroid assay module. Spheroid area was calculated using ImageJ plugin INSIDIA [53].

### Proximity ligation assay (PLA)

PLA was performed on cultured granulosa cells or intact COCs using the Duolink PLA Probes and PLA Fluorescence in situ Detection Kit Red as per manufacturer’s protocol. Briefly, cultured granulosa cells or COCs were fixed in 4% paraformaldehyde and permeabilised with PBS + 0.01% Triton X-100 for 1 h at room temperature. Cells were blocked with blocking Buffer for 1 h at 37°C and incubated with primary antibody couple (N-cadherin + β-catenin) diluted in Antibody Diluent for 2 h at room temperature or overnight at 4°C. Then, cells were incubated with PLA probes of appropriate species for 1 h at 37°C, oligo probes were ligated for 30 min at 37°C and the amplification reaction was carried out at 37°C for a minimum of 100 minutes. Between steps, cells were washed using the provided wash buffers. Slides were mounted with Prolong Gold Mounting Media, cured for at least 1 h in the dark and were stored at −20°C prior to imaging. Slide imaging was done through the Olympus confocal microscope FV1000.

### Isolation of ovaries, cumulus-oocyte complexes and granulosa cells

For the time-course of gene regulation, mice were untreated or were hormonally stimulated by intraperitoneal injection with 5 IU equine chorionic gonadotropin (eCG) followed by 5IU hCG at 46 hour-post eCG. Mice were culled at the following time points: unstimulated (no eCG and hCG), 0 h (no hCG), 4, 8, 10, 12 post-hCG. Ovaries were dissected and granulosa cells were collected by repeated puncturing of the ovaries.

Ovaries or oviducts were dissected from mice and placed in HEPES-buffered minimum essential medium alpha (αMEM) supplemented with 1% (v/v) fetal calf serum at 37°C. Ovulated cumulus-oocyte complexes (COCs) were isolated from the oviducts of mice at 16 h after hCG injection as indicated, placed in HEPES-buffered α-MEM supplemented with 1% (v/v) foetal calf serum and counted under a stereo microscope. After counting, ovulated COCs were snap-frozen for RT-qPCR analyses.

One ovary per mouse was fixed in 4% paraformaldehyde (w/v) in PBS [80 mM Na2HPO4, 20mM NaH2PO4 and 100 mM NaCl (pH 7.5)] for 24 h and processed into paraffin blocks that was then sectioned (5 µm) and stained with Hematoxylin and Eosin. Images were captured at high resolution using NanoZoomer Digital Pathology technology (Hamamatsu Photonics K.K.). The other ovary was snap frozen in liquid nitrogen and stored at −80°C for RNA analysis.

### Gene expression analyses

Total RNA was isolated using Trizol (ThermoFisher Scientific), as per manufacturer’s instructions, with the inclusion of 15 µg GlycoBlue (Ambion) during precipitation. Total RNA was then treated with 1U of DNase as per manufacturer’s instructions. RNA concentration and purity were quantified using a Nanodrop ND-1000 Spectrophotometer (Biolab Ltd., Victoria, Australia). First-strand complementary DNA (cDNA) was synthesized from total RNA using random hexamer primers (Geneworks) and Superscript III reverse transcriptase (Invitrogen). Quantitative reverse transcription PCR (Q-RT-PCR) was performed using Taqman gene expression assays (Applied Biosystems) and reactions were run in duplicate on an AB7900HT Fast PCR System using manufacturers recommended amplification settings. Gene expression levels were normalized to a reference gene (Rpl19).

### Immunofluorescence

Immunohistochemistry and Immunofluorescence were performed on 4% PFA-fixed paraffin-embedded 5-μm sections mounted onto Super frost microscope slides (ThermoFisher Scientific) or whole mounts of denuded oocytes. Tissue sections were dewaxed in xylene and rehydrated. Antigen retrieval was performed by incubating section in either citrate buffer ((10 mM sodium citrate, pH 6.0) or in Tris-EDTA buffer (10 mM Tris/ 1 mM EDTA/ 0.05 % (v/v) Tween 20, pH 9.0) for 20 min at 95°C. After cooling, sections were washed with TBS containing 0.025% Tween-20 (TBST, pH7.6) for 10 mins and blcoked in 10% normal goat serum (NGS) in TBST for 1h at RT. Sections were probed with primary antibodies against N-cadherin (BD Biosciences; Cat# 610920; 1:500); β-catenin (CST; Cat#8480; 1:500), E-cadherin (CST, Cat#14472; 1:500); a-tubulin (ThermoFisher; Cat#236-10501); Ki-67 (CST; Cat#9449; 1:1000) and Cleaved caspase 3 (CST; Cat#9661; 1:1000) diluted in 10% NGS and incubated O/N at at 4°C overnight in a humid chamber. Negative controls were performed using matching isotype control either rabbit IgG (CST; Cat#3900S) or mouse IgG (CST; Cat#5415S) at equivalent concentration used for primary antibodies. Sections were washed three times for 5 min each in TBST and incubated with secondary antibodies Alexa flour 647; Alexa Flour 488; Alexa Flour 594 (ThermoFisher Scientific; Cat# A21244, A11034, A32728; A110001; 1:2000) for 1 h at RT alongside Hoeschst 33342 (ThermoFishe Scientific; Cat#H3570; 1:250). Sections were washed three times for 5 min each in TBST and mouted with flourescence mounting medium (Dako, Santa Clara, USA). Images of immunofluorescence secitons were captured by confocal microcopy FV1000 (Olympus, Tokyo, Japan).

### RNA sequencing and data processing

Total RNA was extracted from COCs and whole ovaries by homogenising in trizol as described above. mRNA was converted to strand specific Illumina compatible sequencing libraries using the Nugen Universal Plus mRNA mRNA-Seq library kit (Tecan, Mannedorf, Switzerland) as per the manufacturer’s instructions. Libraries were sequenced on the Illumina Nextseq550 High output mode and v2.5 chemistry platform for 75bp single-end reads. Sequencing quality control was performed using FastQC. Sequences were mapped to the GENCODE mouse transcriptome (GRCm38.p6, M25) and mapped transcripts were quantified using Salmon. Clustering analysis was performed on count data using variance stabilizing transformation as part of the DESeq2 package. Read count was transformed to log counts per million (CPM) for visualisation via heatmap using edgeR package (R/Bioconductor). Differential expression was analysed using DESeq2. Genes that had a highly stringent FDR <= 10^-6^ and log fold change >=+/- 0.5 were determined to be differentially expressed genes (DEGs). Biological significance of DEGs was explored by GO term enrichment analysis including biological process (BP) and moleuclar function (MF), based on Enrichr web application. Gene Set enrichment analysis of the genes differentially expressed upon CRS-066 treatment was done using a pre-ranked list on the Gene Set Enrichment Analysis software (software.broadinstitute.org/gsea/index.jsp). Gene sets were extracted from the Molecular Signatures database (The Molecular Signatures Database, v7.3) using the search term “β-catenin”, “Hippo”. FDR *q* value was used to rank the results. Gene sets enriched at FDR *q* value ≤ 0.05 and nominal *P*< 0.05 were considered statistically significant. The raw data of all sequencing libraries generated in this study have been submitted to the Gene Expression Omnibus that can be accessed with the number of GSE168347 (ovary) and GSE168348 (COCs).

### Statistical analysis

Statistical analysis was performed using GraphPad Prism 7.0 software (GraphPad Software Inc., La Jolla, CA, USA). Differences between two groups were calculated using the unpaired two-tailed Student’s *t*-test. Statistical analyses of three or more groups were compared using one-way analysis of variance (ANOVA) followed by Bonferroni’s multiple comparisons test. Reported values are the mean ± standard error of the mean of three or more independent biological experiments or as indicated. * p < 0.05 and ** p<0.01.

## Declarations

### Ethics approval and consent to participate

All animal procedures were approved by the University of Adelaide Animal Ethics Committee (Ethics approval IDs M-2018-071 and M-2018-117) and conducted in accordance with the Australian Code of Practice for the Care and Use of Animals for Scientific Purposes.

### Consent for publication

Not Applicable

### Data Availability

The datasets generated during the current study are available in the Gene Expression Omnibus repository, accession ID # GSE168347 and GSE168348

## Funding

This work was supported by grants from the Bill and Melinda Gates Foundation Contraceptive Discovery Program (OPP1171844, INV-001616). DLR is supported by NHMRC Senior Research Fellowship APP1110562. RLR is supported by NHMRC Senior Research Fellowship APP1117976. KM is supported by an Early Career Cancer Research Fellowship from Cancer Council SA’s Beat Cancer Project on behalf of its donors and the State Government of South Australia through the Department of Health and Wellbeing.

### Authors’ contributions

DLR is responsible for conceptualising, funding acquisition, experimental design, and participated in methodology, data analysis and manuscript preparation. AE is responsible for validation and execution of experiments and endpoint methods, data analysis, preparation of figures and manuscript. RLR participated in conceptualising experimental design, and manuscript preparation. DTD and TM contributed methods and data analysis and manuscript editing. OWB provided N-cadherin antagonists and editing of manuscript. RB and AA provided antagonist analogues and manuscript editing. KMM, ACWZ contributed to methods design interpretation and manuscript preparation.

## Acknowledgements

The authors acknowledge the instruments and technical assistance of Microscopy Australia at Adelaide Microscopy, The University of Adelaide.

## Supplementary Figure Legends

**Supplementary figure S1.** Spheroid formation assay in N-cadherin deficient or N-cadherin expressing mouse 67NR cells. (a) Bright-field images of spheroid formation assay in either wild-type 67NR cells lacking endogenous N-cadherin and unable to form spheroids and in 67NRs Cdh2 stably expressing N-cadherin and successfully undergo spheroid compaction within 6 h. (b) Example of automated “spheroid mask” image segmentation used to calculate spheroid area using ImageJ software. (c) Mean +/- SEM of spheroid area in 67NR-Cdh2 or 67NR WT over 6h calculated after “spheroid masking”.

**Supplementary figure S2.** In situ proximity ligation assay (PLA). PLA confirms close-proximity interaction between N-cadherin and β-catenin (red) on granulosa cell junctions (a, b) and at oocyte membrane (c).

**Supplementary figure S3.** (a) Principal component analyses of COCs treated with either CRS-066 (2.5uM) or vehicle (DMSO) for 10h during IVM. (b) Volcano plot of differentially regulated genes (DEGs) (adjusted p<10-6 and log2FC>0.5). (c) Effect of YAP inhibition on cumulus cell expansion and meiotic maturation. (d) Relative mRNA expression of key cumulus expansion genes normalised to Rpl19. Mean and SEM (n=5).

**Supplementary figure S4.** (a) Dosing schedule for in vivo studies with N-Cadherin antagonist CRS-066 (50mg/kg). (b) Principal component analyses of ovaries collected from mice treated with either CRS-066 or vehicle 16h post-hCG. (c) Volcano plot of DEGs (adjusted p<10-6 and log2FC>0.5).

